# Third-generation dual-source computed tomography of the abdomen: comparison of radiation dose and image quality of 80kVp and 80/150kVp with tin filter

**DOI:** 10.1101/2020.03.25.007609

**Authors:** Seung Joon Choi, So Hyun Park, Seong Ho Park, Seong Yong Pak, Jae Won Choi, Su Joa Ahn, Young Sup Shim, Yu Mi Jeong, Bohyun Kim

**Author notes:** Corresponding Author: So Hyun Park, MD., Ph.D., Department of Radiology, Gil Medical Center, Gachon University College of Medicine, 21, Namdong-daero 774beon-gil, Namdong-gu, Incheon, 21565, Korea. Seung Joon Choi and Su Joa Ahn contributed equally to this study.

## Abstract

**Objective:** To compare the radiation dose, objective and subjective image quality, and diagnostic performances of 80 kVp and 80/150 kVp with tin filter (80/Sn150 kVp) computed tomography (CT) in oncology patients.

**Methods:** One hundred forty-five consecutive oncology patients who underwent third-generation dual-source dual-energy CT of the abdomen for evaluation of malignant visceral, peritoneal, extraperitoneal, and bone tumor were retrospectively recruited. Two radiologists independently reviewed each observation in 80 kVp CT and 80/Sn150 kVp CT. Modified line-density profile of the tumor and contrast-to-noise ratio (CNR) were measured. Diagnostic confidence, lesion conspicuity, and subjective image quality were calculated and compared between image sets.

**Results:** Modified line-density profile analysis revealed higher attenuation differences between the tumor and the normal tissue in 80 kVp CT than in 80/Sn150 kVp CT (127 vs. 107, *P =* 0.05). The 80 kVp CT showed increased CNR in the liver (8.0 vs. 7.6) and the aorta (18.9 vs. 16.3) than the 80/Sn150 kVp CT. The 80 kVp CT yielded higher enhancement of organs (4.9 ± 0.2 vs. 4.7 ± 0.4, *P <* 0.001) and lesion conspicuity (4.9 ± 0.3 vs. 4.8 ± 0.5, *P* = 0.035) than the 80/Sn150 kVp CT; overall image quality and confidence index were comparable. The effective dose reduced by 45.2% with 80 kVp CT (2.3 mSv ± 0.9) compared to 80/Sn150 kVp CT (4.1 mSv ± 1.5).

**Conclusions:** The 80 kVp CT performed similar or better than 80/Sn150 kVp CT for abdominal tumor evaluation with 45.2% radiation dose reduction in oncology patients.

## Introduction

Optimization of the radiation dose delivered in abdominopelvic computed tomography (CT) imaging is important, especially in situations where repeated CT examinations are performed in patients. Various strategies have been developed to further reduce the radiation dose, including low kVp, automated exposure control, and iterative reconstruction (IR) [1,2]. Currently, 80 kVp is feasible and increasingly used optimizing radiation dose [3-6]. Although using low kVp reduces the radiation dose, it increases image noise [7]. IR selectively reduces statistical noise in the images, thereby improving the quality of subtle image details and may facilitate dose reduction. In the past decade, evolution from statistical-based to model-based iterative algorithms has improved the performance of IR algorithms [8]. Advanced modelled iterative reconstruction (ADMIRE; Siemens Healthcare, Forchheim, Germany) is a model-based IR that can reduce noise in raw data and may allow further dose reduction while generating images of acceptable quality.

Dual-source dual-energy CT (DECT) systems enable dual-energy data to be acquired using two x-ray sources at different energy levels, i.e., a variety of voltage and tube current combinations [9-12]. Dual-source DECT images using a blend of low- and high-kVp images, provide an image impression similar to a standard 120 kV image (i.e., 80/140, 80/Sn150, or 90/Sn150 kVp), to generate virtual noncontrast images [11]. In other words, low kVp/high kVp with tin filter (Sn) CT generates low kVp, virtual noncontrast, and blended images, which are widely used in oncology [11]. Although initial DECT revealed three times higher radiation dose than single-energy CT [13], recent studies have demonstrated that third-generation dual-source DECTs can be performed without radiation dose penalty or impairment of image quality compared to single-energy CT with 100-120 kVp [14,15]. They suggested that DECT can be routinely used in patients because of no increase in radiation dose compared to single-energy CT. However, no comparison has been reported between images with low kVp from single-energy CT and blending images from DECT. If 80 kVp CT scan performs similar to blending image (80/Sn150 kVp) of DECT while reducing the radiation dose, it may reduce the use of DECT in patients who receive repetitive scans. To our knowledge, this imaging scheme concept has not been explored previously.

Thus, the purpose of our study was to compare the radiation dose and image quality between 80 kVp and 80/Sn150 kVp CT scans performed by the third-generation dual-source DECT using ADMIRE reconstruction algorithm.

## Materials and methods

### Study design

This study is a retrospective analysis of CT images and was approved by our Institutional Review Board with waiver of the requirement for patient consent.

### Study patients

We screened 191 consecutive patients referred to our department between August 2018 and March 2019 for dual-source abdominopelvic CT examination for assessment of response to oncology treatment, or surveillance of known or previously treated malignancy. The inclusion criteria were as follows: (a) solid nodules of malignant liver tumor, other malignant visceral tumor (except liver), metastatic lymph node (larger than 1.5 cm along the short axis), peritoneal tumor, extraperitoneal tumor (i.e., retroperitoneum, muscle, subcutaneous fat layer), and metastatic bone tumor, (b) the nodules ≤ 5 in number, and (c) nodules not typically hemangiomas or cysts. Six predetermined abdominopelvic lesions modified from the previously described report [16]. We limited the number of nodules to minimize cluster bias. We excluded 46 patients without a reference standard (MRI, PET/CT, or surgery), lack of a lesion on CT, or because of a change in protocol. Thus, 259 nodules in 145 consecutive patients were included (Fig. 1). Nodules were selected and annotated by an experienced study coordinator (BLINDED-FOR-PEER-REVIEW), who was not involved in the image analysis.

**Fig. 1.**
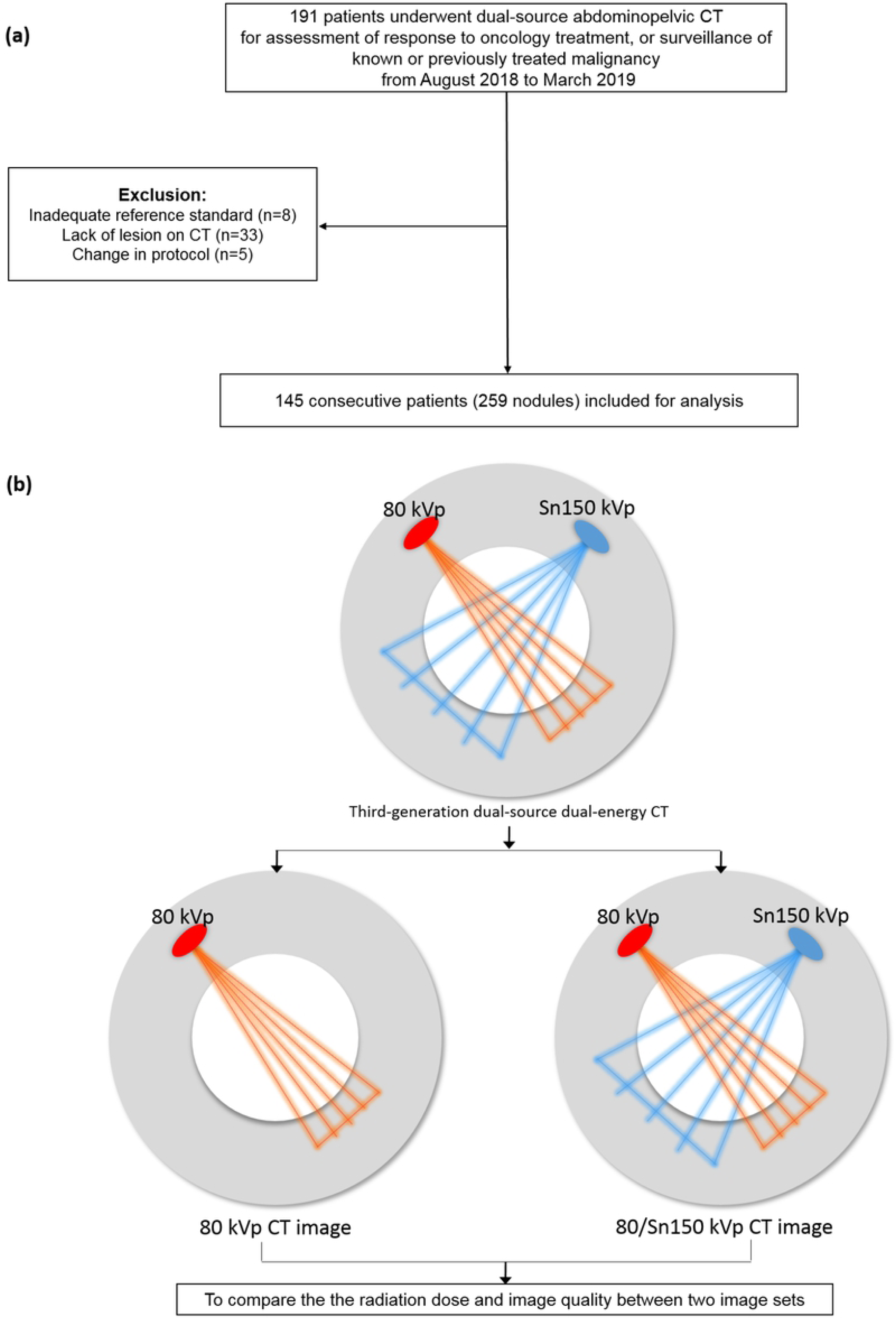
Patients’ flow chart including selection process (a) and study design (b)

### CT examination protocol

All CT scans were performed using a 192-slice, third-generation dual-source CT scanner (SOMATOM Force, Siemens Healthcare, Forchheim, Germany) in the dual-energy mode with tube detector A (80 kVp; reference tube current, 250 mAs) and B (Sn 150 kVp; reference tube current, 125 mAs), using tube current dose modulation (CARE dose 4D; Siemens Healthcare) (Table 1).

**Table 1.** CT acquisition scanning parameters

|  |  |  |
| --- | --- | --- |
|  | 80 kVp | 80/Sn150 kVp |
| Detector tube | A tube | Combination of<br>A and B tube |
| Kilovolt | 80 | 80/Sn150 |
| Tube current time product (mAs) | 250 | 250/125 |
| Coverage |  |  |
| Acquisition mode | Helical | Helical |
| z-flying focal spot | 192 × 0.6 mm | 192 × 0.6 mm |
| Automated tube current modulation | On | On |
| Automated tube voltage selection | Off | Off |
| Pitch | 1.15 | 1.15 |
| Rotation time (sec) | 0.5 | 0.5 |
| Reconstruction algorithm | ADMIRE | ADMIRE |
| Thickness of axial image | 3 mm | 3 mm |

Note. ADMIRE, advanced modelled iterative reconstruction.

### Data reconstruction

Images from 80/Sn150 kVp CT (raw data from both tube detectors) and 80 kVp CT (raw data from tube detectors A) scans were reconstructed using the ADMIRE algorithm at a strength level of 2 out of 5 with axial slice thicknesses of 3 mm each with edge-enhancing convolution kernel (Br64). The reconstruction was performed on a multimodality workstation (syngo.via VB20, Siemens Healthcare). The image was reviewed on our clinical picture archiving and communication system (PACS, INFINITT Healthcare, Korea).

### CT data analysis

#### Subjective analysis: image quality and diagnostic confidence

Interpretations of 290 examinations were performed by two independent board-certified radiologists (BLINDED-FOR-PEER-REVIEW, with 10 and 12 years of experience in abdominopelvic CT) in two reading sessions of 80 kVp and 80/Sn150 kVp CT scans. Each reading session included half of the two image sets, with a 6-week washout period between sessions to avoid recall bias. The images were reviewed anonymously, with the order of review randomized separately for each review session. After each session, consensus results between readers were used to resolve discrepancies. Overall image quality for diagnostic purpose was graded using a 5-point scale (1, nondiagnostic image quality, strong artefacts; 2, severe artefacts with uncertainty about the evaluation; 3, moderate artefacts with mild restricted assessment; 4, slight artefacts with unrestricted diagnostic image evaluation possible; and 5, excellent image quality with no artefacts). Enhancement of organs was graded using a 5-point scale (1, very poor; 2, suboptimal; 3, acceptable; 4, above average; and 5, excellent). Image noise was graded using a 5-point scale (1, unacceptably high; 2, higher than average; 3, average; 4, less than average; and 5, minimum noise). A score was derived for axial images with a soft-tissue window setting (width, 400 HU; level, 50 HU) on a PACS system. Reviewers were allowed to change the window level and width as per their comfort level during analysis.

For each presented acquisition, both readers had to independently assess the predetermined abdominopelvic lesions. Each item was graded using a five-point confidence index (1, very poor; 2, poor; 3, average; 4, high; and 5, excellent confidence) and an ordinal scale for lesion conspicuity (1, definite artefact [pseudolesion]; 2, probable artefact; 3, subtle lesion; 4, poorly visualized margins; and 5, well-visualized margins). A score of either 1 or 2 was considered unacceptable for diagnostic purposes.

#### Objective image analysis

Image noise, defined as standard deviations (SDs) in Hounsfield units (HU), were measured by drawing a circular region of interest (ROI; size between 1 and 3 cm^2^) on axial images of three image sets in the following three anatomical regions by one blinded reader (BLINDED-FOR-PEER-REVIEW) [17-19]: the subcutaneous fat in the anterior abdominal wall, the lumen of abdominal aorta, and the right hepatic lobe parenchyma. Mean attenuation values were measured (in HU) for each ROI in all three image sets. Signal-to-noise ratio (SNR = mean HU of interested area/fat noise) and contrast-to-noise ratio (CNR = [mean HU of interested area - mean HU of psoas muscle]/ fat noise) were calculated [7,16]. Fat noise was calculated as the mean of standard deviation of CT attenuation in ROIs placed within the mesenteric fat. ROIs were carefully placed at the same location between different image series.

The two image sets were used for modified line-density analysis [20]. The tumor was identified on 80/Sn150 images, and one reader, with 4 years of experience, positioned a line of 10 mm in length and 2 mm in width perpendicular to the tumor margins with one half of the line within the tumor and the other half within the normal tissue. Mean, minimum, and maximum HU values were measured within this 10-mm line using a multimodality workstation. Modified line-density analysis was performed on 128 nodules from 92 patients, as not enough normal was available in patients with peritoneal seeding lesions.

### Radiation dose

The volume CT Dose Index (CTDI_vol_) and dose-length product (DLP) were recorded from the scanner dose page. The effective dose (in millisieverts, mSv) was calculated using the tissue-weighting factor for the abdomen (male, *k* = 0.013; female, *k* = 0.017) and pelvis (male, *k* = 0.010; female, *k* = 0.016) based on international commission on radiological protection publication 103 with modification [21,22] (an average value; male, *k* = 0.012; female, *k* = 0.017). Size-specific dose estimates (SSDEs) were calculated by measuring the effective diameter from the anteroposterior and lateral dimensions on the CT scan. A conversion factor based on the effective diameter and a 32-cm-diameter polymethylmethacrylate phantom were used [23].

### Statistical analysis

Dose parameters and image analysis were compared between different dose CT scans using Student’s t-test and chi-squared test. The interobserver agreement between the two radiologists for each of the assessed subjective image quality parameter was estimated using intraclass correlation coefficients (ICCs) and applying the following scale: 0.01–0.20, slight; 0.21–0.40, fair; 0.41–0.60, moderate; 0.61–0.80, substantial; and 0.81–1, excellent agreement.

Interobserver agreement regarding image features was estimated using the overall proportion of agreement [24], rather than κ statistics, as the latter is affected by the distribution of data across categories, thus, not a valid indicator of our data [25]. A *p*-value < 0.05 was considered statistically significant. All statistical analyses were performed using SPSS Statistics for Windows, Version 21.0 (IBM Corp.).

## Results

### Patient characteristics

Of the 145 patients in our study, 79 (54.5%) were men and 66 (45.5%) were women with mean age ± standard deviation of 62.4 ± 10.5 years. The body mass index (BMI) ranged from 15.1 to 30.7 kg/m^2^ (mean, 22.8 kg/m^2^; median, 22.5 kg/m^2^). Altogether, 259 nodules were included in the analysis: 98 malignant nodules in the liver, 55 peritoneal seeding nodules, 45 metastatic bone tumors, 31 metastatic lymph nodes, 23 other visceral tumors, and 7 extraperitoneal tumors (Table 2).

**Table 2.** Patient information

| Parameter | Value |
| --- | --- |
| **Patients, (n = 145)** |  |
| Age (years) * |  |
| All patients (men: women = 79: 66) | 62.4 ± 10.5 |
| Height (cm) * | 163.1 ± 8.2 |
| Weight (kg) * | 60.2 ± 12.6 |
| Effective diameter (cm) * | 26.2 ± 2.9 |
| Body mass index (BMI) * | 22.8 ± 4.0 |
| <18.5 (underweight) | 23 (15.9) |
| 18.5–24.9 (normal) | 81 (55.9) |
| 25–29.9 (overweight) | 33 (22.8) |
| 30–34.9 (moderately obese) | 6 (4.1) |
| 35–39.9 (severely obese) | 2 (1.4) |
| Type of primary cancer |  |
| Colorectal | 47 (32.4) |
| Hepatobiliary | 25 (17.2) |
| Gastro-small bowel | 24 (16.5) |
| Breast | 17 (11.7) |
| Genitourinary | 14 (9.7) |
| Bronchial | 8 (5.5) |
| Others | 10 (6.9) |
| Reference standard |  |
| Liver MRI and PET/CT | 48 (33.1) |
| PET/CT | 37 (25.5) |
| Pelvis MRI and PET/CT | 18 (12.4) |
| PET/CT and spine MRI | 17 (11.7) |
| Liver MRI, PET/CT, and surgery | 15 (10.3) |
| Liver MRI, PET/CT, and spine MRI | 7 (4.8) |
| Liver MRI and surgery | 3 (2.0) |

Note. Data are presented as number (%), unless indicated otherwise.
\* Data are means ± standard deviations.

MRI, magnetic resonance imaging; PET, positron emission tomography; CT, computed tomography

### Objective image quality

The 80 kVp CT had significantly higher attenuation compared to 80/Sn150 kVp CT in the liver and aorta (125.9 HU vs. 107.2 HU, *P <* 0.001; 208.2 HU vs. 160.5 HU, *P <* 0.001; Table 3; Fig. 2). The CNR of the aorta in 80 kVp CT images was significantly higher than in 80/Sn150 kVp CT images (*P <* 0.001) whereas the CNR of the liver in 80 kVp CT images was mildly higher than that in 80/Sn150 kVp CT images (*P* = 0.197). The SNR of the liver and aorta tended to be higher in 80/Sn150 kVp CT images, although there was no significant difference between the image sets (*P* = 0.075 and 0.069, respectively).

**Table 3.**
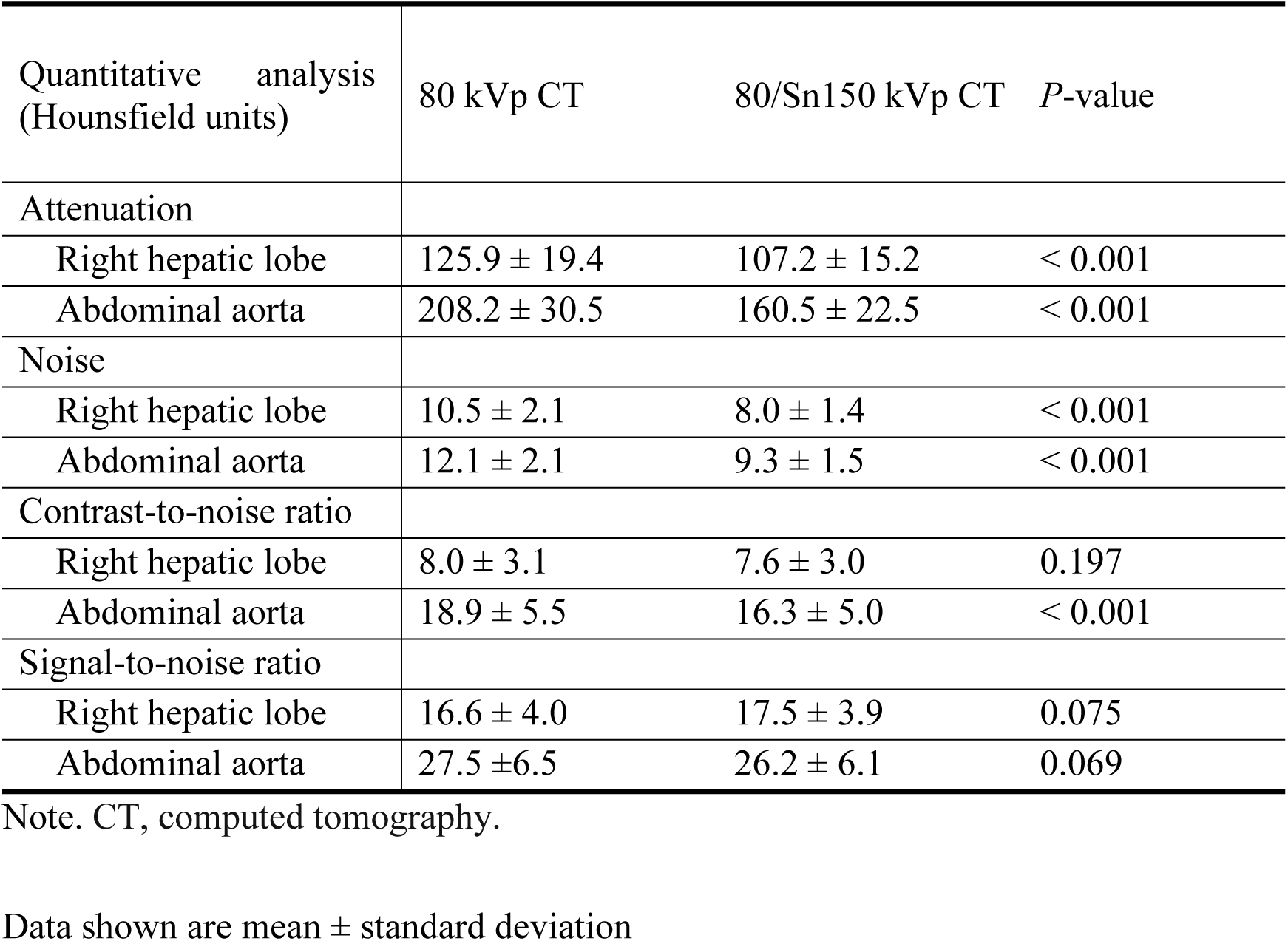
Assessment of objective image quality

| Quantitative analysis<br>(Hounsfield units) | 80 kVp CT | 80/Sn150 kVp CT | P-value |
| --- | --- | --- | --- |
| Attenuation |  |  |  |
| Right hepatic lobe | 125.9 ± 19.4 | 107.2 ± 15.2 | < 0.001 |
| Abdominal aorta | 208.2 ± 30.5 | 160.5 ± 22.5 | < 0.001 |
| Noise |  |  |  |
| Right hepatic lobe | 10.5 ± 2.1 | 8.0 ± 1.4 | < 0.001 |
| Abdominal aorta | 12.1 ± 2.1 | 9.3 ± 1.5 | < 0.001 |
| Contrast-to-noise ratio |  |  |  |
| Right hepatic lobe | 8.0 ± 3.1 | 7.6 ± 3.0 | 0.197 |
| Abdominal aorta | 18.9 ± 5.5 | 16.3 ± 5.0 | < 0.001 |
| Signal-to-noise ratio |  |  |  |
| Right hepatic lobe | 16.6 ± 4.0 | 17.5 ± 3.9 | 0.075 |
| Abdominal aorta | 27.5 ± 6.5 | 26.2 ± 6.1 | 0.069 |

Note. CT, computed tomography.
Data shown are mean ± standard deviation

**Fig. 2.**
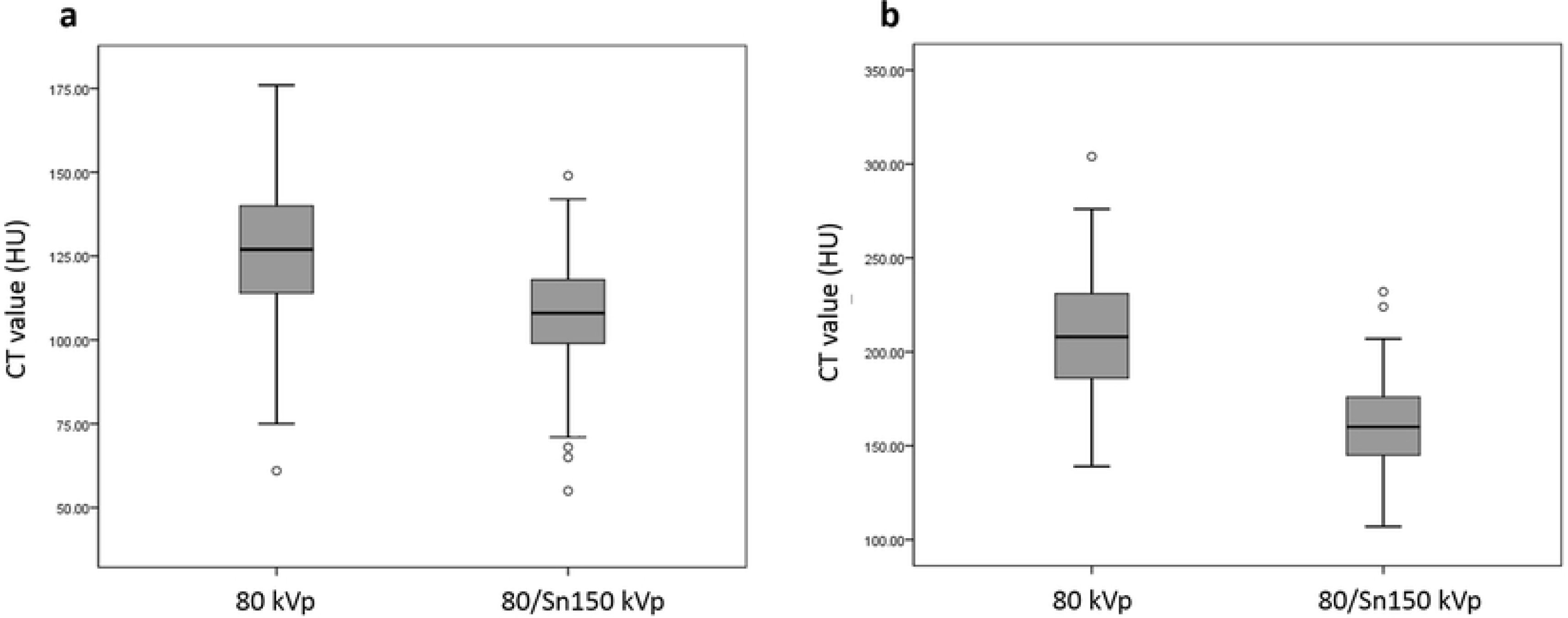
Box and whisker plots of CT enhancement in abdominal aorta (a) and liver parenchyma (b) between 80 kVp and 80/Sn150 kVp. Box-and-whisker diagrams show mean values ± standard deviation, as well as minimum and maximum values.

The mean differences between maximum and minimum attenuation within the tumor are shown in Table 4. The mean attenuation differences within the tumor in 80 kVp CT were significantly higher than those in 80/Sn150 kVp CT (127.2 HU vs. 107.0 HU, *P =* 0.05).

**Table 4.** Differences between modified line-density profiles within the tumor border

| Hounsfield units | 80 kVp | 80/Sn150 kVp | P-value |
| --- | --- | --- | --- |
| Mean ± SD | 127.2 ± 71.9 | 107.0 ± 59.4 | 0.05 |
| Maximum | 882 | 726 |  |
| Minimum | -66 | -54 |  |

Note. The difference was defined as the maximum Hounsfield units – minimum Hounsfield units. SD, standard deviation

### Subjective image quality

The 80 kVp CT showed significantly higher enhancement of organs (4.9 ± 0.2 vs. 4.7 ± 0.4, *P <* 0.001) and lesion conspicuity (4.9 ± 0.3 vs. 4.8 ± 0.5, *P* = 0.035) than 80/Sn150 kVp CT according to the consensus interpretation and similar overall image quality (4.8 ± 0.3 vs. 4.9 ± 0.2) and confidence index (4.8 ± 0.5 vs. 4.8 ± 0.6; Fig. 3-4, Table 5).

**Table 5.** Assessment of subjective image quality.

|  | 80 kVp CT | 80/Sn150 kVp CT | *P*-value |
| --- | --- | --- | --- |
| Overall image quality | 4.8 ± 0.3 | 4.9 ± 0.2 | 0.306 |
| Enhancement of organs | 4.9 ± 0.2 | 4.7 ± 0.4 | < 0.001 |
| Image noise | 4.8 ± 0.3 | 4.9 ± 0.1 | 0.061 |
| Confidence index | 4.8 ± 0.5 | 4.8 ± 0.6 | 0.687 |
| Lesion conspicuity | 4.9 ± 0.3 | 4.8 ± 0.5 | 0.035 |

Note. CT, computed tomography.
Data shown are mean ± standard deviation

**Fig. 3.**
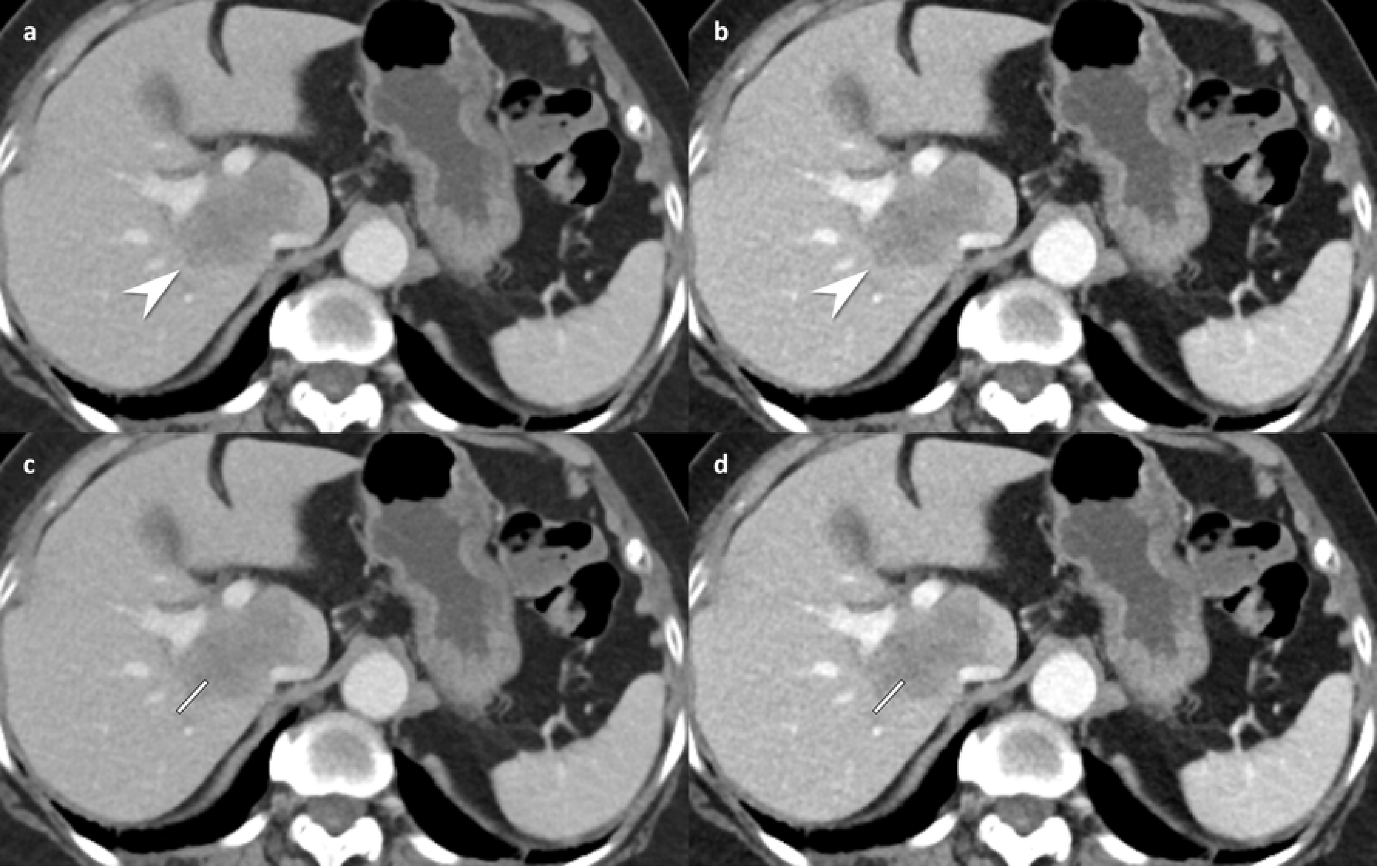
Contrast-enhanced abdomen CT in portal venous phase in a 37-year-old woman (body mass index, 29.0 kg/m^2^) with ascending colon cancer and liver metastasis. 80/Sn150 kVp CT (a, effective dose; 5.9 mSv) shows a liver metastasis in the right hepatic lobe/segment 1 (white arrowhead) by each reader (reader1 and reader2, grade 4 lesion conspicuity, grade 5 diagnostic confidence, and grade 5 overall image quality). Corresponding image obtained during the same contrast-enhanced phase with 80 kVp CT (b, effective dose; 3.4 mSv) demonstrates increased conspicuity of the lesion by each reader (white arrowhead, reader1 and reader2, grade 5 lesion conspicuity, grade 5 diagnostic confidence, and grade 5 overall image quality). Modified line-density profile revealing higher attenuation differences between the tumor and the normal tissue at 80 kVp (d) than those at 80/Sn150 CT (c) (maximum HU: minimum HU, 157:52 vs.129:49). CT, computed tomography.

**Fig. 4.**
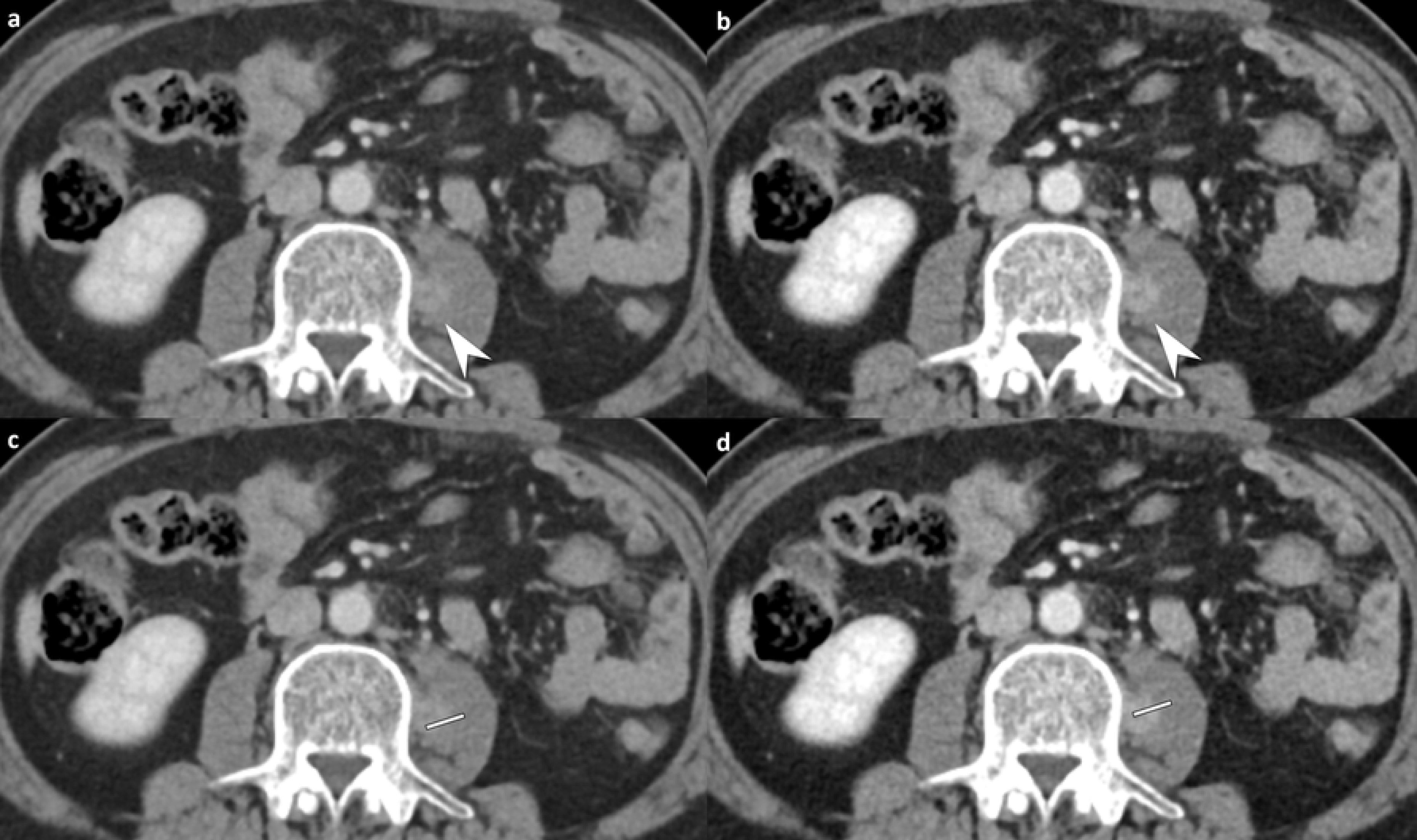
Contrast-enhanced abdomen CT in portal venous phase in a 45-year-old man (body mass index, 24.8 kg/m^2^) with sigmoid colon cancer with a seeding nodule in the left psoas muscle. 80/Sn150 kVp CT (a-b, effective dose; 4.5 mSv) showing a hyperattenuating mass in the left psoas muscle (white arrows) by each reader (reader1 and reader2, grade 4 lesion conspicuity, grade 5 diagnostic confidence, and grade 5 overall image quality). The corresponding image obtained during the identical phase with 80 kVp CT (effective dose; 2.6 mSv) demonstrates markedly increased conspicuity of the lesion in the left psoas muscle by each reader (reader1 and reader2, grade 5 lesion conspicuity, grade 5 diagnostic confidence, and grade 5 overall image quality). Modified line-density profile revealing higher attenuation differences between the tumor and the normal tissue at 80 kVp (d) than those at 80/Sn150 CT (c) (maximum HU: minimum HU, 129:27 vs.106:36). CT, computed tomography.

#### Reader agreement

Interobserver agreement between the two readers for the overall image quality, enhancement of organs, image noise, confidence index, and lesion conspicuity were 85.5%, 92.0%, 87.2%, 89.1%, and 84.3 %, respectively.

#### Radiation dose

Various dose parameters of the two image sets are shown in Table 6 and Figures 5. For the 80/Sn150 kVp CT, the mean CTDI_vol_ was 5.3 ± 1.4 mGy, and the SSDE was 7.4 ± 3.8 mGy with an effective dose of 4.1 ± 1.5 mSv. For the 80 kVp CT, the mean CTDI_vol_ was 2.9 ± 1.0 mGy, and the SSDE was 4.1 ± 2.2 mGy with an effective dose of 2.3 ± 0.9 mSv. On average, the effective dose of 80 kVp CT scan was 45.2% ± 2.8% (range, 38.0–51.9%) less than that of 80/Sn150 kVp CT.

**Table 6.**
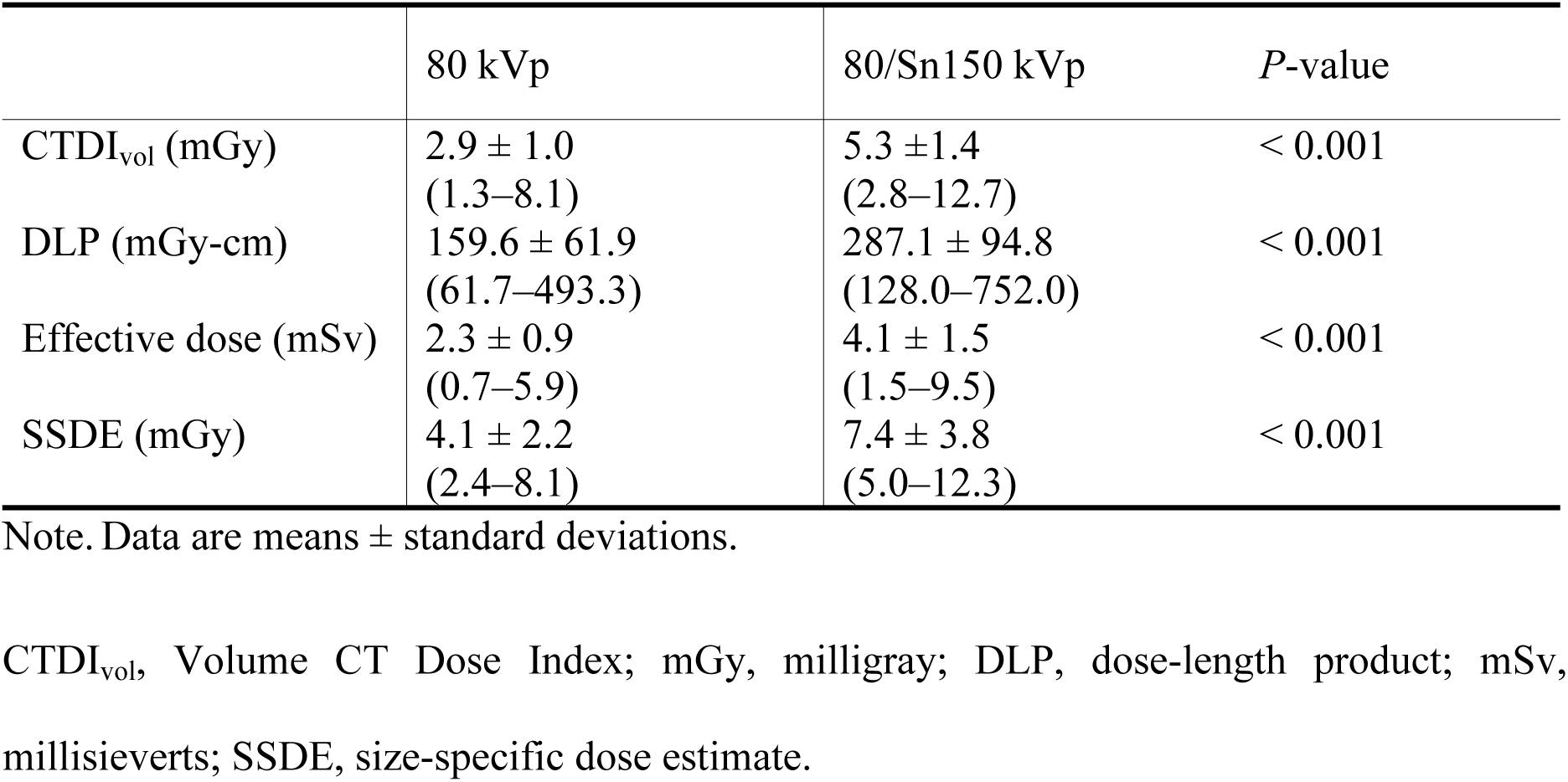
Dose parameters of each image set

|  | 80 kVp | 80/Sn150 kVp | *P*-value |
| --- | --- | --- | --- |
| CTDI<sub>vol</sub> (mGy) | 2.9 ± 1.0<br>(1.3–8.1) | 5.3 ± 1.4<br>(2.8–12.7) | < 0.001 |
| DLP (mGy-cm) | 159.6 ± 61.9<br>(61.7–493.3) | 287.1 ± 94.8<br>(128.0–752.0) | < 0.001 |
| Effective dose (mSv) | 2.3 ± 0.9<br>(0.7–5.9) | 4.1 ± 1.5<br>(1.5–9.5) | < 0.001 |
| SSDE (mGy) | 4.1 ± 2.2<br>(2.4–8.1) | 7.4 ± 3.8<br>(5.0–12.3) | < 0.001 |

Note. Data are means ± standard deviations.
CTDI<sub>vol</sub>, Volume CT Dose Index; mGy, milligray; DLP, dose-length product; mSv, millisieverts; SSDE, size-specific dose estimate.

**Fig. 5.**
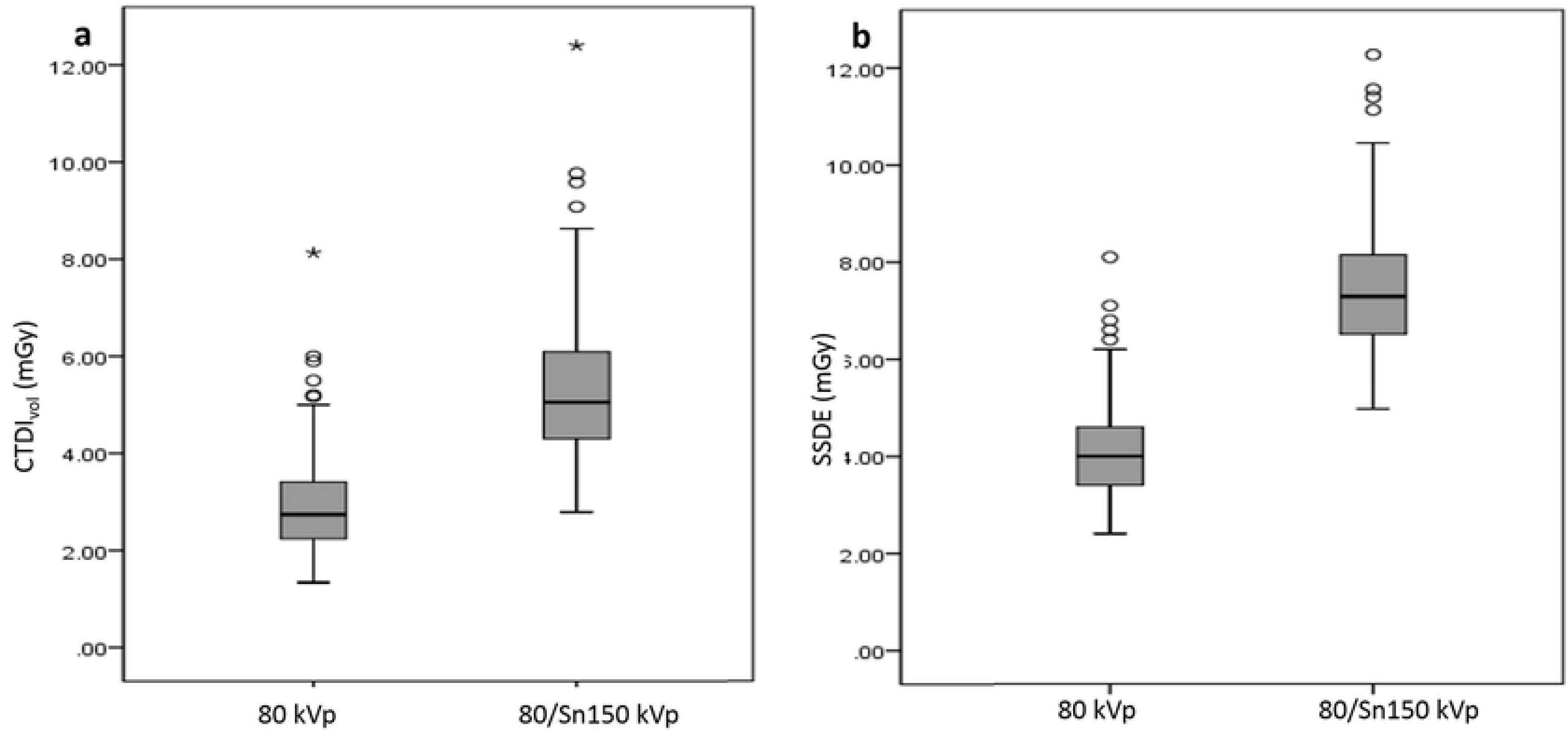
Box and whisker plots of CTDI_vol_ (a) and SSDE (b) for 80 kVp and 80/Sn150 kVp scans. Box-and-whisker diagrams show mean values ± standard deviation, as well as minimum and maximum values.

## Discussion

We investigated whether the performance of 80 kVp CT was comparable to that of the 80/Sn150 kVp CT while reducing the radiation dose, to ascertain if it could be used in oncology patients who receive repetitive DECT scans. Our findings revealed that there was a 45.2% reduction in radiation dose by using the low tube voltage (80 kVp) CT imaging with ADMIRE while maintaining excellent image quality compared with DECT (80/Sn150 kVp). Contrast-related features including enhancement of organs, CNR, and attenuation differences within the tumor border were higher in 80 kVp CT than in 80/Sn150 kVp CT, while SNR showed no significant differences. A potential explanation for these findings is that lowering the CT tube voltage results in a greater photoelectric effect of iodinated contrast media, with only a slight increase in the noise because of improved tube current capabilities; moreover, the advances in hardware equipped with Stellar detectors and software program in third-generation dual-source CT [26,27], which are supposed to be more sensitive to electron influx are thus, dose-efficient and generate high quality images while using 80 kVp. A few studies that investigated DECT, showed that the radiation dose was comparable to the single energy 100-120 kVp CT [14,28]. However, our study found that the 80 kVp CT showed superior image quality with lower radiation dose than DECT in abdominal scans. Thus, DECT should not be routinely used in patients who may undergo abdominal CT repeatedly.

Several studies have reported that 80 kVp abdominal CT can be used effectively while reducing radiation dose in patients with normal BMI [29-32]. Studies have also reported that reducing the tube voltage from 120 kVp to 80 kVp resulted in a 48-65% dose reduction using either an identical tube current or automated tube current modulation [33-35]. However, extremely low tube voltage scans may result in noisier images that are inadequate for interpretation. Increase in tube current or use of IR may improve image quality and lesion conspicuity [26]. Our study showed that the effective dose of 80 kVp CT (2.3 mSv ± 0.9) scan was 45.2% less than 80/Sn150 kVp CT (4.1 mSv ± 1.5) using IR. Low tube-voltage CT scan is a robust method for radiation dose reduction in abdominal CT.

Third-generation dual-source CT is more dose-efficient than second-generation dual-source CT because of technical advances and adjusted scan protocols while the image quality is consistently high with all assessed protocols [32]. The low kVp is more likely selected while using automated tube voltage modulation (ATVM), thereby reducing the radiation dose in third-generation dual-source CT. Part et al. reported that third generation dual-source CT demonstrated an increase in the number of 90-100 kVp CTs whereas the majority of the patients were selected on 100-120 kVp using ATVM on second generation dual-source CT [32]. Winklehner et al. reported that using third-generation dual-source CT compared to second-generation, resulted in a tube voltage decrease of at least 10 kV in most patients (75%, 100 kVp→90 kVp) who received body CT angiography examinations [35]. These results support the use of low kVp abdominal CT in patients for reducing radiation dose as well as for maintaining the diagnostic performance employed in our study.

Low tube-voltage CT technique improves tumor conspicuity and tumor-to-tissue CNR [3,4]. In our study, mean differences between the maximum and minimum attenuation within the tumor was 127.2 HU in 80 kVp CT and 107.0 HU in 80/Sn150 kVp CT. Our results also demonstrated that 80 kVp CT scan might be preferred over 80/Sn150 kVp CT for detecting hypervascular tumors because of better scores in all contrast-related features, increased enhancement of organs, CNR, and attenuation differences within the tumor border. The lesion conspicuity was slightly higher in 80 kVp CT compared to 80/Sn150 kVp CT, which means low-density lesion detection, is superior in 80 kVp CT compared to 80/Sn150 kVp CT. However, SNR of the liver and the aorta showed no significant difference between the image sets in our study. Our results showed a balance between image noise and low-density lesion detectability while using low tube-voltage CT to evaluate solid organs.

DECT in abdomen has the advantage of analyzing CT images in various ways, including monoenergetic image analysis, liver fat and iron quantification, urinary calculi characterization, and gallstone imaging [12,33,34]. Unlike previous studies [14,28] that suggested routine use of DECT, our study found that the performance alone was not sufficient to justify routine use of DECT for oncology patients who frequently undergo CT examinations, sometimes every 1-2 months, because the radiation dose is higher with 80/Sn150 kVp CT compared 80 kVp CT. Moreover, we reported superior tissue contrast features with 80 kVp CT.

Our study had some limitations. First, it was a retrospective study involving oncology patients with tumor lesion on abdominal CT, which may have introduced selection bias. Second, our study had a small number of patients with moderate to severe obesity; thus, we could not perform a robust comparison between 80/Sn150kVp and 80kVp CT in patients with moderate to severe obesity. Third, 80/Sn150kVp scan combines 80kVp and Sn150kVp to generate the image, therefore, 80/Sn150kVp was set to have an unconditional higher radiation dose than 80kVp scan. However, our aim was to focus on evaluating diagnostic performance and image quality between low kVp image using single-energy scan and blended image generated using DECT scan. Rather than increasing the radiation exposure of the patient through two scans, we thought that the intraindividual comparison would help ethically as well. Finally, we only evaluated the performance of a single low kVp image and a blended image. However, various combinations of sequences can build distinctive protocols of 80/140kVp, 90/Sn150kVp, 90/Sn150kVp compared with 70-100 kVp CT scan.

In conclusion, 80 kVp CT showed a decrease in radiation dose exposure, and superior or comparable objective and subjective image quality, compared to DECT. The use of 80 kVp CT using ADMIRE can safely reduce the radiation dose in a patient when compared with 80/Sn150 kVp CT.

## ABBREVIATIONS

ADMIRE: advanced Modeled Iterative Reconstruction
ATVM: automated tube voltage modulation
CNR: contrast-to-noise ratio
CTDI_vol_: volumetric CT dose index
DLP: dose length product
DECT: dual-energy computed tomography
ED: Effective dose
HU: Hounsfield units
IR: iterative reconstruction
mGy: milligray
mSv: millisieverts
ROI: region of interest
SD: standard deviation
SNR: signal-to-noise ratio
SSDE: size-specific dose estimate

